# Putative origin of *Myxococcus fulvus* 124B02 plasmid, pMF1, from a chromosomal segment in another *Myxococcus* species

**DOI:** 10.1101/2024.04.09.588723

**Authors:** Shruti Jain, Gaurav Sharma

## Abstract

Myxobacteria or order Myxococcales (old nomenclature) or phylum Myxococcota (new terminology) are fascinating organisms well known for their diverse peculiar physiological, taxonomic, and genomic properties. Researchers have long sought to identify plasmids within these organisms, yet thus far, only two organisms from different families have been found to harbor a plasmid. This study delves into the origin and evolution of one of these plasmids, i.e., pMF1 present in *M. fulvus* 124B02 in the suborder Cystobacterineae and family Myxococcaceae. Here we first reannotated the pMF1 plasmid genome sequence and identified two additional genes which were not annotated till now. We further reported that all pMF1 plasmid genes depict homology with *M. stipitatus* CYD1 draft genome (contig 28) and a chromosomal segment of *M. stipitatus* DSM14675 in a syntenic manner, suggesting the presence of plasmid-like structure in *M. stipitatus* CYD1 but integrated as part of its chromosome. To comprehend the relationship among these three species, we conducted phylogenetic analyses using 16S and concatenated housekeeping genes and GGDC analysis, which confirmed that *M. stipitatus* CYD1 is a distinct and novel species within the genus *Myxococcus*. Overall, this study sheds light on the putative origin of the pMF1 plasmid, which likely originated from a closely related yet distinct species, possibly through the partition from its chromosome as a segment.

**Importance:** Myxobacteria are not well-known to have plasmids. Till now, only two organisms have been shown to have plasmids, raising a pertinent question about how these plasmids evolved randomly within the phylum Myxococcota. The study presented in this manuscript delves into the origin and evolution of the pMF1 plasmid found in *Myxococcus fulvus* 124B02, a member of the suborder Cystobacterineae and family Myxococcaceae. Our research addresses the intriguing topic of plasmid identification and evolution within Myxobacteria, which are a group of fascinating organisms that have garnered significant interest due to their diverse physiological, taxonomic, and genomic properties.

## Introduction

Microorganisms are well known for being “ecosystem suppliers” as they are crucial for the environment and human life (Johansen, 2021). Microbes have adapted themselves hugely according to their respective niches via diverse processes such as gene duplication, genome rearrangement, horizontal gene transfer, gene loss, etc. (Feero et al., 2011). Myxobacteria, order Myxococcales (or phylum Myxococcota as of recent ICSP nomenclature) members, are gram-negative bacteria well known for their extraordinary multicellular social lifestyle. Their peculiar characteristics are influenced by their multicellular behavior, such as predation, cooperative movement, or social gliding motility (Kaiser et al., 2010) and multicellular structures known as fruiting bodies (Muñoz-Dorado et al., 2016). 16S rRNA studies have shown that the myxobacteria group belongs to the class Deltaproteobacteria (Wrótniak-Drzewiecka et al., 2016), and the majority of these organisms are aerobic except for facultative anaerobe *Anaeromyxobacter* spp. and strict anaerobe *Pajaroellobacter abortibovis*. Together with actinomycetes, *Bacillus* spp., and several fungi, myxobacteria are among the vast producers of diverse natural products such as polyketides, non-ribosomal polypeptides, terpenoids, phenylpropanoids, alkaloids, etc. Many of these compounds are effective against bacteria, viruses, fungi, cancer cells, and other microorganisms. Genetic engineering can play a pivotal role in exploiting the potential of myxobacterial secondary metabolites by utilizing their plasmids. Till 2000, various myxobacteria species were screened, but no indigenous plasmid was found in any species. To re-evaluate whether myxobacterial organisms contain plasmids or not, about 150 strains belonging to the genus *Myxococcus* and a few genera *Corallococcus* (phylogenetically closely related genus of family Myxococcaceae and suborder Cystobacterineae) were rescreened, and a circular plasmid pMF1 was discovered in *M. fulvus* strain 124B02 (Zhao et al., 2008). In 2021, another plasmid, pSa001, was identified from *Sandaracinus sp*. MSr10575, which belongs to the family Sandaracinaceae and suborder Sorangiineae (Panter et al., 2021). However, pMF1 remains the only plasmid known for suborder Cystobacterineae and genus Myxococcus, the most studied model organism in the order Myxococcales.

*M. fulvus* strain 124B02 is an aerobic, gram-negative bacteria having slender rod-shaped vegetative cells with tapering ends and typical *Myxococcus*-type fruiting bodies containing spherical myxospores. *M. fulvus* 124B02 (*Mf*124B02) genome consists of a circular chromosome with a total length of 11.04 Mbp, 69.96% GC content, and 8,515 predicted coding sequences (Chen et al., 2016). Three completely identical 16S rRNA sequences are present in its genome in three different 16S-23S-5S operons located at 0.24, 7.35, and 10.84 Mb sites.

Plasmid pMF1 is a low copy number plasmid with a genome size of 18,634 bp. Its GC content (68.7%) is very similar to the Mf124B02 chromosome sequence (Zhao et al., 2008). Twenty-three ORFs have been predicted for this plasmid, of which 21 are on the sense strand and two (pMF1.19c and pMF1.20c) on the complementary strand. About 86% of the plasmid sequence consists of protein-coding sequences, although the function of 14 ORFs is mainly unknown (Chen et al., 2016). The region pMF1.13-pMF1.16 is the replication origin locus, out of which pMF1.14 is essential for plasmid propagation (Zhao et al., 2008). pMF1.19-pMF1.20 region functions like a post-segregational killing system to improve plasmid maintenance (Li et al., 2018). ORFs pMF1.19-pMF1.20 were previously described as SitAI toxin-immunity pairs where the SitA toxin is delivered by the TraAB outer membrane exchange system (Vassallo & Wall, 2019, PNAS). pMF1.21-pMF1.23 region helps retain plasmids’ stability (Sheng et al., 2021; Sun et al., 2011). Till now, no apparent beneficial genes, such as genes coding for antibiotic resistance, virulence, or growth factors, have been identified, which are required for its persistence in the host (Chen et al., 2016).

Research thus far has provided insights regarding replication locus, partitioning system, and post-segregational killing system in this plasmid. However, as the functions of most of the plasmid genes are not known, further studies are required to comprehend what particular advantage this plasmid provides to this bacterium and how it originated evolutionarily. Here we investigated in-depth annotations of this plasmid and figured out how this plasmid originated in this organism or broadly within myxobacteria.

## Methodology

### Data collection

The complete genome (GCA_000988565.1) and plasmid pMF1 (EU137666.1) information for *Mf*124B02 were retrieved from the National Center for Biotechnology Information (NCBI) GenBank database (Benson et al., 2018). The Plasmid pMF1 genome was reannotated using RAST (Rapid Annotations with Subsystem Technology) (Overbeek et al., 2014) and Prokka (version 1.14.6) (Seemann, 2014) to certify that no other genes are present in the plasmid. The location of genes annotated from RAST, Prokka, and NCBI was compared to determine the unique number of genes in the plasmid **(Table-S1)**. The final translated CDS file for the plasmid containing all the annotated genes was used for further exploration. A genome statistics table of *Mf*124B02 complete genome, plasmid (NCBI annotated), and merged plasmid annotation based on genome size, total number of genes, coding density, CDS (positive strand), CDS (negative strand), GC content, etc. was prepared **(Table-1)**.

### Function prediction of all annotated genes and understanding the origin of plasmid pMF1

The input plasmid genome sequence was subjected to PLSDB, an open-source plasmid database (Schmartz et al., 2022), to find its homology with available plasmid sequences. To remove the possibility of plasmid’s sporadic or continuous homology within the *Mf*124B02 genome, plasmid pMf1 gene/protein sequences were subjected to standalone blastn/blastp (version 2.11.0) against *Mf*124B02 gene/protein sequences. To decipher the relationship of the plasmid with other organisms, blastp analysis (Johnson et al., 2008) for all plasmid-translated CDS sequences was performed against the NCBI non-redundant (NR) database. Taxonomy information was obtained for each blast hit using the “efetch” tool and parsed with the result. Homologs and their corresponding taxonomy information of all hits were analyzed to determine if there is any synteny between the pMF1 plasmid and other organisms. *M. stipitatus CYD1 (MsCYD1)* and *M. stipitatus DSM 14675 (MsDSM14675)* translated CDS files were downloaded from the NCBI RefSeq database, and the individual database for both files was generated using makeblastdb. Standalone blastp (version 2.11.0) was performed to compare the plasmid pMF1 protein sequences against the *Ms*CYD1 and *Ms*DSM14675.

### Genome annotation of *Ms*CYD1 and *Ms*DSM14675

Contig 28 of *Ms*CYD1 was reannotated using RAST and Prokka (version 1.14.6) and the locations of all annotated genes using RAST, Prokka and NCBI were compared followed by the identification of new genes. All pMF1 translated proteins were subjected to blastp against *Ms*CYD1 proteins with 1e^-10^ evalue cutoff. Remaining plasmid and *Ms*CYD1 genes that didn’t show the homology were compared using Clustal Omega. Similarly, the remaining plasmid and *Ms*DSM14675 genes were also compared using Clustal Omega. Finally, a synteny diagram was generated using ggplot2 and geom_gene_arrow method of gggenes library in R to represent blastp alignment results among the pMF1, *Ms*CYD1 and *Ms*DSM14675.

### Comparative genomics of genus *Myxococcus* spp

Genome sequences and associated files of 53 *Myxococcus* species and an outgroup, *Corallococcus coralloides* DSM2259, were downloaded from NCBI and a genome statistics list with information about genome size, GC content, total contigs, total genes, and total RNAs number were tabulated **(Table-S2)**. To understand the evolutionary relationship of *Mf*124B02, *Ms*CYD1 and *Ms*DSM14675 among all 53 *Myxococcus* spp., 16S rRNA sequences with the longest length per organism were extracted. Subsequently, standalone muscle (version 3.8.1551) (Edgar, 2004) was used to run the multiple sequence alignment. Model selection for phylogeny was performed using megacc (version 7.0) (Tamura et al., 2021) and then RAxML (version 8.2.12) (Stamatakis, n.d.) was used to build the phylogeny using the maximum likelihood method with identified GTRGAMMA model and 100 bootstrap values. RAxML bipartitions file was processed to generate the newick file, which was further visualized in the Interactive Tree of Life (iTOL) tool (Letunic & Bork, 2021). A list of thirty-one housekeeping genes was used to mine homologous housekeeping genes in 53 *Myxococcus* species and 1 *Corallococcus* species, using standalone blastp. Thirty-one housekeeping genes were identified and extracted in all 54 organisms. Each gene set was subjected to multiple sequence alignment using standalone MUSCLE (version 3.8.1551) followed by sorting by headers and concatenated to create one final alignment file. The final alignment was subsequently subjected to model selection using megacc (version 7.0) followed by building the phylogeny using RAxML (maximum likelihood method, PROTGAMMALG model, 100 bootstraps). Genome to Genome Distance Calculator (GGDC) was used to calculate the distance between all 54 genomes, thereby facilitating genome-to-genome comparison (Meier-Kolthoff et al., 2022). The *Mf*124B02 genomic file was used as the query sequence and the reference sequence (against which the genome comparison is performed) was a set of all 53 *Myxococcus* organisms. To unravel the distance of *Ms*CYD1 with other *Myxococcus* species, GGDC was also run with MsCYD1 as a query.

## Results and Discussion

### Genome reannotation of pMF1

*Mf*124B02 has a complete genome assembly of 10.784 Mb with 90.46% coding density, whereas the plasmid pMF1 genome is 18.634 Kb long with 85.6% coding density. NCBI annotation suggests 23 genes, of which two genes are present on the negative strand and the remaining 21 on the positive strand. As a single annotation might outlook a few genes, the pMF1 genome was reannotated using RAST (Overbeek et al., 2014) and Prokka (Seemann, 2014), and both predicted 25 ORFs. A combined annotation was generated using homology, gene frame, and location comparative of all ORFs from all annotations, suggesting the number of final annotated genes in all possible combinations **(Figure-S1; Table-S1)**. Out of 23 NCBI annotated genes, 15 genes were found homologous to the other two annotations, whereas out of 25 Prokka and RAST annotated genes, 21 and 19 genes were homologous to the other two annotations, respectively. Several gene combinations were partial overlaps, and the longest ORF was finally selected amongst such homologs. This study revealed that only one Prokka gene overlaps with the RAST annotation, but no RAST gene was exact homolog with the Prokka annotation. Based on overlapping and homology analysis, we found 1, 4, and 5 genes uniquely annotated by Prokka, RAST, and NCBI, respectively. Based on a comparative merging of all three (NCBI, RAST, and Prokka) annotations, 25 genes were finalized **(Table-1)**, which includes two new genes, i.e., located at 1-744 (new1), and 5540-6103 bp (new2) on the positive strand. **(Table-S1)**. This reannotation-based collated information increased the plasmid coding density from 85.6% to 92.6 **(Table-1)**.

### Deciphering origin of plasmid based on homology analysis

Blast analysis against **the** PLSDB (Schmartz et al., 2022) database indicated that the plasmid pMF1 has no homology with other known existing plasmids. We observed only self-hits for all 25 pMF1 genes (Figure-S2). To confirm that this plasmid has not originated from its host, plasmid gene/protein sequences were subjected to blastp-based homology analysis against *Mf*124B02 complete genome. The results portrayed that only 4 genes out of 25 genes show homology with the complete genome, therefore, ruling out the possibility that some portion of complete genome might have duplicated and separated as an independent plasmid (Figure-S2). To find any relevant homology with any available sequence, blast analysis (Johnson et al., 2008) of plasmid sequences against the NR database revealed no significant hits for new1, new2 and ABX46789.1_6 whereas 8 genes namely ABX46792.1_3, ABX46792.1_9, ABX46797.1_14, ABX46798.1_15, ABX46800.1_17, ABX46801.1_18, ABX46804.1_21 and ABX46806.1_23 showed only self-hit. In this analysis, the remaining 14 genes showed homologs in either *Ms*CYD1 or *Ms*DSM14675 genomes.

### Homology analysis of plasmid genes suggests their putative evolution from *M. stipitatus* CYD1 via circularization of a DNA fragment

To confirm the homology and putative origin from *M. stipitatus* strains, a standalone blast was specifically performed against both *M. stipitatus* strains, which revealed that out of 25 plasmid pMF1 genes, 22 and 11 genes show good identity with *Ms*CYD1 (draft genome) and *Ms*DSM14675 (complete genome) genes, respectively. These 22 and 11 genes cover the whole plasmid and approximately half of the plasmid, as depicted in **Figure-1**. All 22 identified *Ms*CYD1 genes are present on a single contig (contig-28; NZ_JAKCFI010000028.1) in a syntenic manner. This contig was further reannotated to identify additional genes in between. According to NCBI annotation, 19 genes were present in contig 28, whereas a hybrid annotation using RAST, Prokka, and NCBI annotation resulted in 23 genes. Intriguingly ABX46784.1_1 gene (second gene of plasmid) shows similarity with prot_3038 (first gene of the contig) as well as with prot_WP_234072500.1_3056 (last gene of the contig) (Table S3). Overall, we can stipulate that this *Ms*CYD1 contig might have circularized to form a plasmid which might have transferred in *Mf*124B02 during evolution.

**Figure-1:**
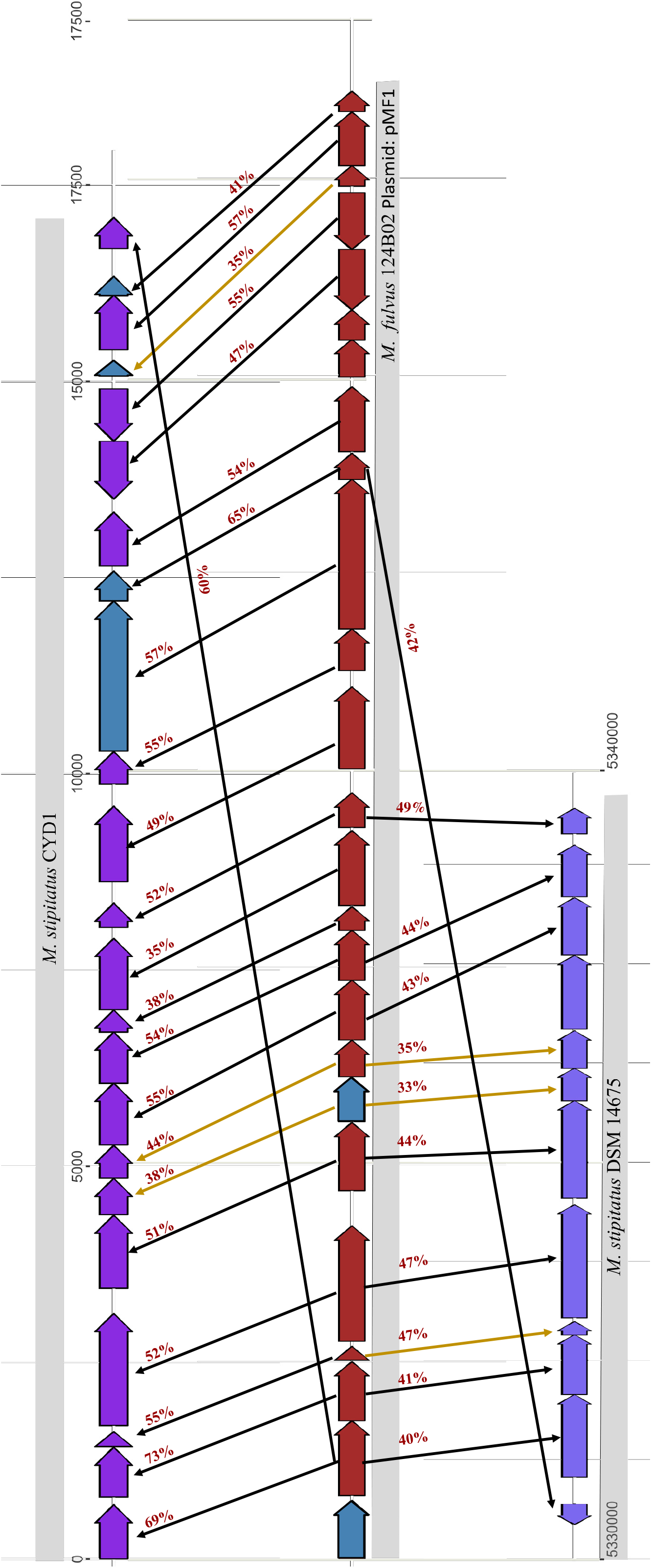
Comparative homology amongst the genes of plasmid *M. fulvus 124B02* pMF1 (Brick red), *M. stipitatus* CYD1 (Dark Purple), and *M. stipitatus* DSM14675 (Light Purple) depicting syntenic relationship along with percentage identity. Blue color boxes are the newly annotated genes in the pMF1 and *Ms*CYD1 genomes. Black and Yellow color lines represent the percentage amino acid identity obtained using BLASTp and Clustal Omega respectively.

All 11 homologs in *Ms*DSM14675 are also present linearly, starting from the prot_WP_015349713.1_4096 gene to prot_WP_015349724.1_4107. Here, the first plasmid gene (ABX46784.1_1) shows similarity with prot_WP_015349714.1_4097 (second gene of that specific region in DSM14675) and seventeenth plasmid gene (ABX46798.1_15) shows similarity with prot_WP_015349713.1_4096 gene (first gene of that particular region of DSM14675) indicating partial similarity with *Ms*DSM14675 genome **(Table-S3)**. Standalone blast analysis of *Ms*CYD1 genes against *Ms*DSM14675 revealed that all these eleven DSM14675 genes also show similarity with *Ms*CYD1 genes, and these CYD1 genes showed similarity with the same plasmid genes. *Ms*CYD1 is a draft genome and has almost all homologs of the plasmid pMF1; however, *Ms*DSM14675, even being a complete genome and close relative of *Ms*CYD1, have only a few homologs portraying that although there are some similarities between *Ms*CYD1 and *Ms*DSM14675 in terms of this DNA segment, *Ms*CYD1 genes might have contributed to the origin and evolution of this plasmid.

Further, we want to comprehend how similar are MsCYD1 and MsDSM14675 to each other, and whether MsCYD1 strain is similar to Mf124B02. Therefore, to answer these questions, we performed comparative genomic studies amongst the available genus Myxococcus organisms.

### Phylogenetic analysis and GGDC reveal *M. stipitatus* CYD1 and *M. fulvus* 124B02 to be distinct species

Fifty-four available genomes (53 *Myxococcus* and one outgroup, i.e., *Corallococcus coralloides* DSM2259) were selected for this study. *Myxococcus* spp. have genome size ranging from 8.8 Mb (*Myxococcus sp. AM009*) to 12.41 Mb (*M. llanfairpwllgwyngyllgogerychwyrndrobwllllantysiliogogogochensis* AM401), GC content ranging from 68 to 70% and total RNA content (including rRNA, tRNA, ncRNA and other RNA) ranging from 68 to 95. 16S rRNA sequences of all organisms whose size ranges from 1100-1600 bp were extracted. *Corallococcus coralloides* DSM2259 was chosen as an outgroup as it belongs to the same family as *Myxococcus*. The reliability of this 16S rRNA phylogenetic tree can be confirmed as most of the *M. xanthus*, and *M. fulvus* are present in their respective clades **(Figure-2)**. It is evident from the analysis that *Mf*124B02 and *Ms*CYD1 are not closely related as both are present in different clades. It further supports that *Ms*CYD1 and *Mf*124B02 are two different organisms. Although 16S rRNA-based phylogeny is popular and helpful, its credibility in associating taxonomic relationships between organisms below the genus level is always dubious (Janda & Abbott, 2007). We also observed one such doubtful issue from this analysis: irrespective of belonging to the same species, *Ms*CYD1 and *Ms*DSM14675 are not present in the same clade. To understand it precisely, we opted for housekeeping genes-based phylogenetic analysis followed by genome-genome distance calculations. Thirty-one conserved housekeeping genes (dnaG, frr, infC, nusA, pgK, pyrG, rplA, rplB, rplC, rplD, rplE, rplF, rplK, rplL, rplM, rplN, rplP, rplS, rplT, rpmA, rpoB, rpsB, rpsC, rpsE, rpsI, rpsJ, rpsK, rpsM, rpsS, smpB and tsf) were used to build a concatenated genes phylogeny. This analysis revealed that all *M. fulvus* strains are closely related to each other as they all belong to the same clade (**Figure-3**).

**Figure-2:**
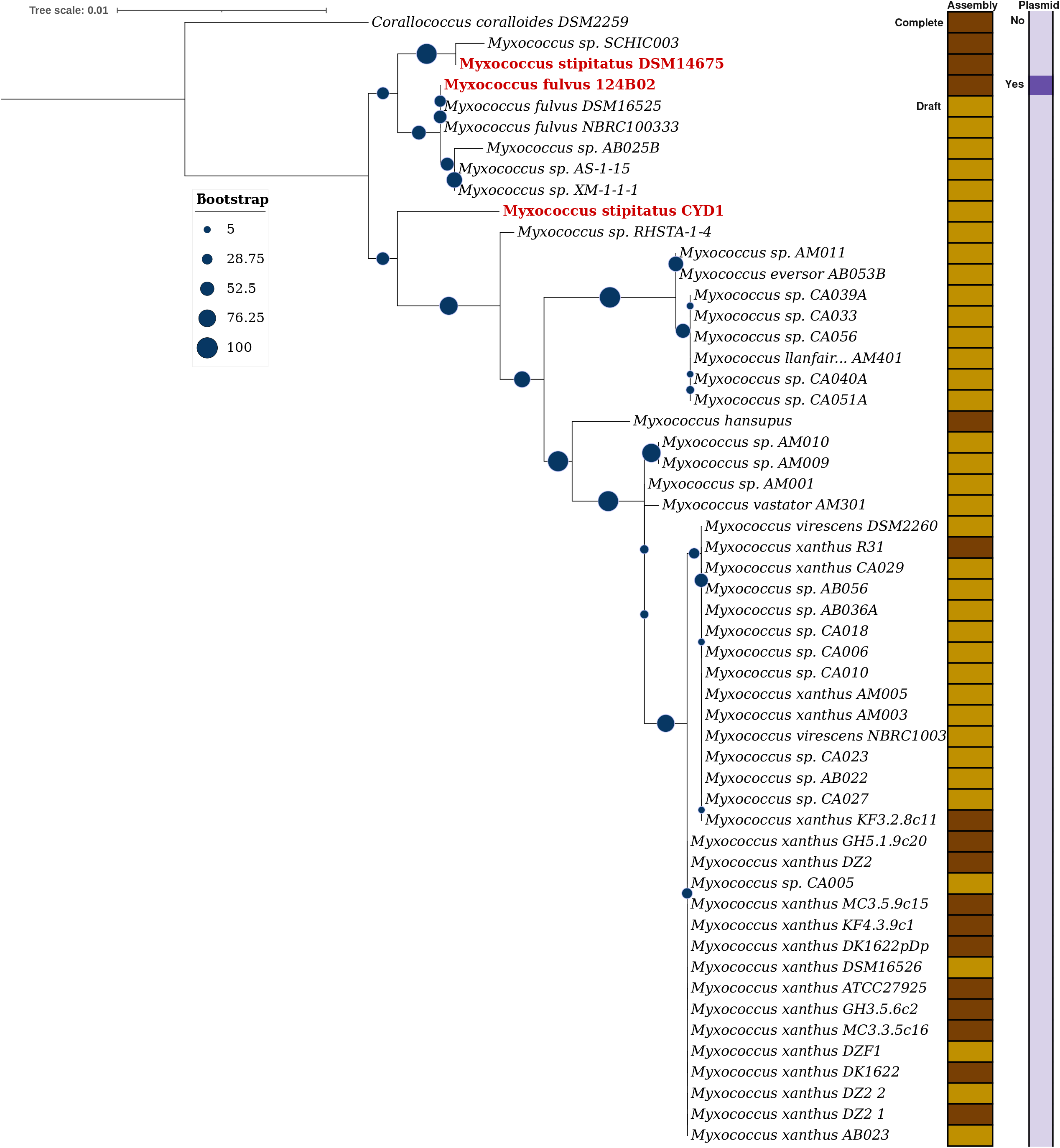
Phylogenetic tree constructed using genus *Myxococcus* spp.16S rRNA sequences and visualized using iTOL. *Corallococcus coralloides* DSM2259 is used as an outgroup. The right-side panels depict the assembly level (complete/draft) and the presence (yes/no) of the plasmid in that organism.

**Figure-3:**
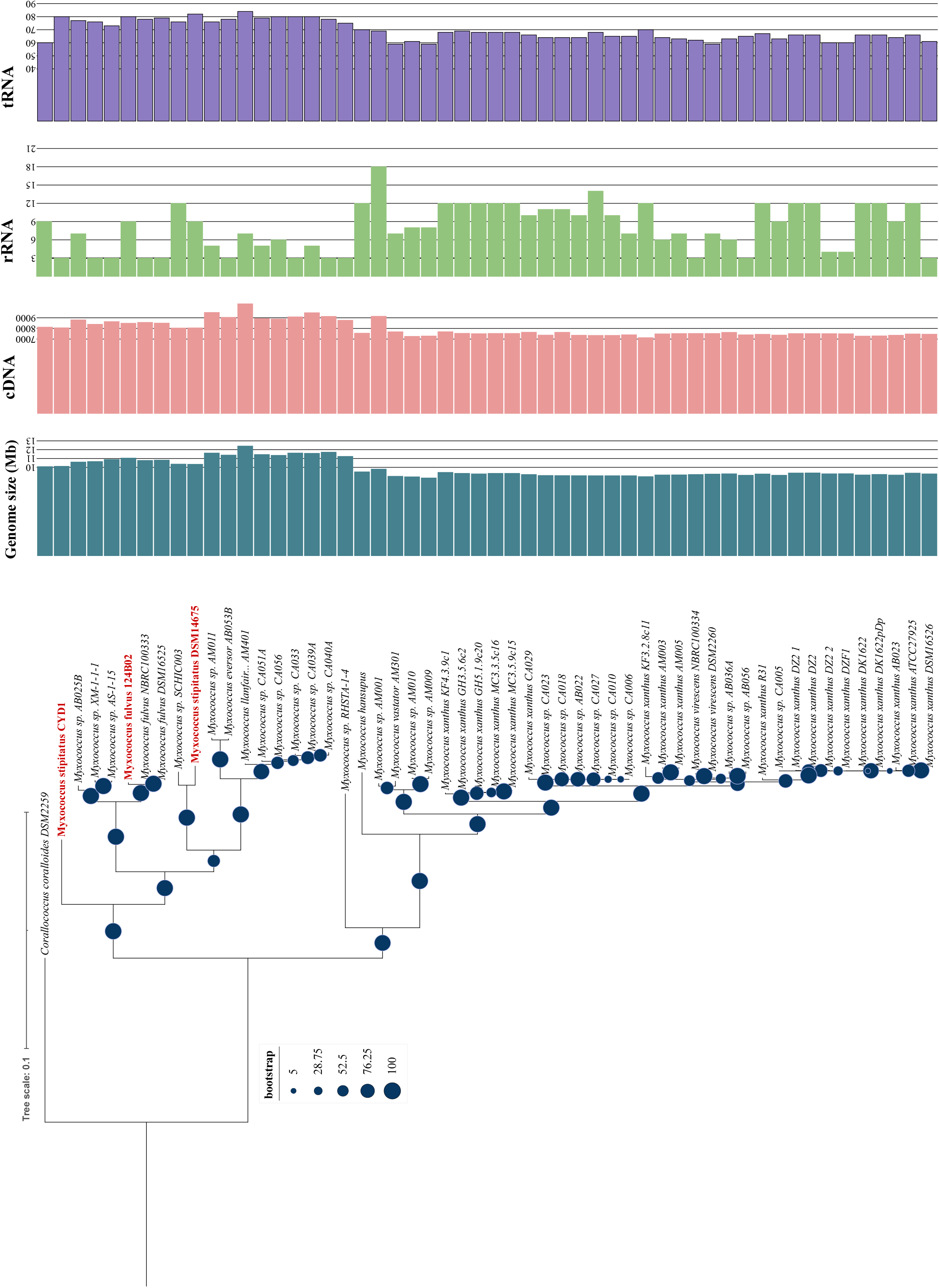
31 Housekeeping genes-based phylogenetic tree: This tree has been visualized in iTOL. Right side panels: cyan color bar plot depicts genome size, pink color bar plot shows cDNA counts, green color plot represents rRNA counts, and purple color plot represents tRNA counts per organism.

*M. xanthus* and *M. virescens* are known to be similar to each other; therefore, most of the strains belonging to these two species are present in their respective clade. This study further supported that *Mf*124B02 and *Ms*CYD1 are not closely related as they belong to different clades. In concordance with 16S phylogeny, *Ms*CYD1 and *Ms*DSM14675 belong to different clades suggesting that *Ms*CYD1 might be a distinct organism compared to *M. stipitatus* and *M. fulvus*. To prove this further, we used **a** genome-to-genome distance calculator as a tool. A DDH (DNA-DNA hybridization) value of >70 usually suggests the query organism to be of the same species. When we looked for the DDH calculations for *Mf*124B02 against all 53 *Myxococcus* spp., we found that all *M. fulvus* spp. have >70 DDH values suggesting their belonging to the same species, whereas the DDH value was 26.5 and 25.3 against *Ms*DSM14675 and *Ms*CYD1 respectively indicating that *Mf*124B02 is distantly related with *M. stipitatus*. For the *MsCYD1* query genome, the DDH value was 24.6 (distance: 0.1773) against *MsDSM14675*, validating that MsCYD1 might be a novel species compared to *M. stipitatus* **(Table-S4)**. *Ms*CYD1 is still in the draft stage and not characterized properly; therefore, we will request culture collection researchers and other experimental biologists to confirm further if *Ms*DSM14675 and *Ms*CYD1 share similar characteristics. If not, then *Ms*CYD1 must be classified as a novel species compared to *Ms*DSM14675.

Overall, based on this analysis, we can speculate two theories in terms of pMF1 origin and evolution: 1) If complete genome sequencing of *Ms*CYD1 reveals that contig 28 is a part of chromosome, which is highly likely as a syntenic homologous segment is present in *Ms*DSM14675, it will suggest that that an ancestral segment related to contig 28 became separated, circularized, and transferred to *Mf*124B02 as a plasmid in its course of time. However, 2) if complete genome sequencing of *Ms*CYD1 reveals that contig 28 is not part of a complete chromosome assembly whereas is a separate plasmid DNA segment, it will suggest that *Ms*CYD1 also contains a plasmid, which might have been transferred to an ancestor of *Mf124*B02. At present, owing to the presence of homologous syntenic segments in *Ms*CYD1, *Ms*DSM14675, and pMF1, the possibility of the first hypothesis is more. Complete genome sequencing and experimental microbial characterization will reveal more in-depth insights. We would like to mention that recently another plasmid-bearing genome, *Myxococcus sp*. MxC21-1, was sequenced and assembled (Liu et al., 2023); however, we did not find good similarity of any gene within this organism as the evalue for best homolog for each pMF1 gene is less than 1E^-5^ (Table S5).

## Conclusion

Overall, this study first reannotated the pMF1 plasmid genome finding two additional genes, further reporting that all pMF1 plasmid genes show syntenic homology within a contig of *M. stipitatus* CYD1 genome (contig 28). This evidence suggests that this *M. fulvus* 124B02 plasmid might have evolved from an *M. stipitatus* CYD1 genomic fragment. This segment is an apparent part of the chromosome as around 11 out of 25 genes are also present in the complete genome of *Ms*DSM14675, which is further supported by the fact that no plasmid-specific genes are present on it. Using 16S and housekeeping genes phylogeny and GGDC analysis, we compared all *Myxococcus* spp. to understand the relationship amongst *Ms*DSM14675, *Ms*CYD1, and *Mf*124B02. This study further confirmed that M. stipitatus CYD1 is a distinct and novel species within the genus Myxococcus, and it should not be considered a member of *M. stipitatus*. Overall, this study discusses the solitary occurrence of a plasmid in myxobacteria showcasing its origin from a chromosomal segment in a close-related but distinct species.

## Figure legends

**<u>Table-1</u>:** Genome statistics of *M. fulvus* 124B02 complete genome, plasmid pMF1, and reannotated plasmid pMF1.

## Supplementary Data Legends

**<u>Figure-S1</u>**: Venn Diagram depicting the comparative gene annotations via Prokka, RAST, and NCBI.

**<u>Figure-S2</u>**: Number of plasmid homologous sequences against plasmid database (PLSDB), *M. fulvus* 124B02 complete genome, and NR database

**<u>Table-S1</u>**: Comparative locations of all annotated genes predicted by Prokka, Rast, and NCBI. For each gene, gene tag EU137666.1_prot_ABX46784.1_1 depicts its genome id followed by prot tag and then protein id.

**<u>Table-S2</u>**: Genome statistics of all studied genus *Myxococcus* spp. and *Corallococcus coralloides* DSM2259 (outgroup) genomes.

**<u>Table-S3</u>**: Homology analysis of plasmid pMF1 genes against *M. stipitatus* CYD1 and *M. stipitatus* DSM 14675

**<u>Table-S4</u>**: GGDC results showing the distance between *M. fulvus* 124B02 and other *Myxococcus* organisms and *M. stipitatus* CYD1 and other *Myxococcus* organisms.

**<u>Table-S5</u>**: Homology analysis of plasmid pMF1 proteins against *Myxococcus sp*. MxC21.

## Ethics approval and consent to participate

Not applicable

## Availability of data and materials

Authors have used open-source tools in this analysis. All tool versions have been provided in the methodology.

## Authors’ contributions

GS generated the idea. SJ performed the whole computational analysis and wrote the first draft of the manuscript. SJ and GS edited and finalized the manuscript.

## References

1. Al Doghaither, H., & Gull, M. (2019). Plasmids as Genetic Tools and Their Applications in Ecology and Evolution. In Plasmid. IntechOpen. 10.5772/intechopen.85705

2. Auch, A. F., Klenk, H. P., & Göker, M. (2010). Standard operating procedure for calculating genome-to-genome distances based on high-scoring segment pairs. Standards in Genomic Sciences, 2(1), 142–148. 10.4056/sigs.541628

3. Beceiro, A., Tomás, M., & Bou, G. (2013). Antimicrobial resistance and virulence: A successful or deleterious association in the bacterial world? In Clinical Microbiology Reviews (Vol. 26, Issue 2, pp. 185–230). 10.1128/CMR.00059-12

4. Benson, D. A., Cavanaugh, M., Clark, K., Karsch-Mizrachi, I., Ostell, J., Pruitt, K. D., & Sayers, E. W. (2018). GenBank. Nucleic Acids Research, 46(D1), D41–D47. 10.1093/nar/gkx1094

5. Bertani, B., & Ruiz, N. (2018). Function and Biogenesis of Lipopolysaccharides. EcoSal Plus, 8(1). 10.1128/ecosalplus.esp-0001-2018

6. Cabeen, M. T., & Jacobs-Wagner, C. (2005). Bacterial cell shape. In Nature Reviews Microbiology (Vol. 3, Issue 8, pp. 601–610). 10.1038/nrmicro1205

7. Carroll, A. C., & Wong, A. (2018). Plasmid Persistence: Costs, Benefits and the Plasmid Paradox 5 6 1125 Colonel By Drive Ottawa, ON K1S 5B6. In Can. J. Microbiol. Downloaded from http://www.nrcresearchpress.com by UNIVERSITY OF CONNECTICUT on. http://www.nrcresearchpress.com

8. Chen, X. jing, Han, K., Feng, J., Zhuo, L., Li, Y. jie, & Li, Y. zhong. (2016). The complete genome sequence and analysis of a plasmid-bearing myxobacterial strain Myxococcus fulvus 124B02 (M 206081). Standards in Genomic Sciences, 11(1), 1–9. 10.1186/s40793-015-0121-y

9. Eberhard, W. (1989). REVIEW Why Do Bacterial Plasmids Carry Some Genes and Not Others? In PLASMID (Vol. 21).

10. Edgar, R. C. (2004). MUSCLE: Multiple sequence alignment with high accuracy and high throughput. Nucleic Acids Research, 32(5), 1792–1797. 10.1093/nar/gkh340

11. F. Panter, C. D. Bader, R. Müller (2021). The Sandarazols are Cryptic and Structurally Unique Plasmid-Encoded Toxins from a Rare Myxobacterium. Angew. Chem. Int. Ed. 2021, 60, 8081

12. Feero, W. G., Guttmacher, A. E., & Relman, D. A. (2011). Genomic Medicine Microbial Genomics and Infectious Diseases. In n engl j med (Vol. 365).

13. Harrison, P. W., Lower, R. P. J., Kim, N. K. D., & Young, J. P. W. (2010). Introducing the bacterial “chromid”: Not a chromosome, not a plasmid. Trends in Microbiology, 18(4), 141–148. 10.1016/j.tim.2009.12.010

14. Janda, J. M., & Abbott, S. L. (2007). 16S rRNA gene sequencing for bacterial identification in the diagnostic laboratory: Pluses, perils, and pitfalls. In Journal of Clinical Microbiology (Vol. 45, Issue 9, pp. 2761–2764). 10.1128/JCM.01228-07

15. Johnson, M., Zaretskaya, I., Raytselis, Y., Merezhuk, Y., McGinnis, S., & Madden, T. L. (2008). NCBI BLAST: a better web interface. Nucleic Acids Research, 36(Web Server issue). 10.1093/nar/gkn201

16. Kaiser, D., Robinson, M., & Kroos, L. (2010). Myxobacteria, polarity, and multicellular morphogenesis. In Cold Spring Harbor perspectives in biology (Vol. 2, Issue 8). 10.1101/cshperspect.a000380

17. Letunic, I., & Bork, P. (2021). Interactive tree of life (iTOL) v5: An online tool for phylogenetic tree display and annotation. Nucleic Acids Research, 49(W1), W293–W296. 10.1093/nar/gkab301

18. Li, Y. J., Liu, Y., Zhang, Z., Chen, X. J., Gong, Y., & Li, Y. Z. (2018). A postsegregational killing mechanism for maintaining plasmid PMF1 in its Myxococcus fulvus host. Frontiers in Cellular and Infection Microbiology, 8(AUG). 10.3389/fcimb.2018.00274

19. Liu L, Xu F, Lei J, Wang P, Zhang L, Wang J, Zhao J, Mao D, Ye X, Huang Y, Hu G, Cui Z and Li Z (2023). Genome analysis of a plasmid-bearing myxobacterim Myxococcus sp. strain MxC21 with salt-tolerant property. Front. Microbiol. 14:1250602. doi: 10.3389/fmicb.2023.1250602

20. Masterson, R. V, Russell, P. R., & Atherly, A. G. (1982). Nitrogen Fixation (nij) Genes and Large Plasmids of Rhizobium japonicum. In JOURNAL OF BACTERIOLOGY. https://journals.asm.org/journal/jb

21. Meier-Kolthoff, J. P., Carbasse, J. S., Peinado-Olarte, R. L., & Göker, M. (2022). TYGS and LPSN: A database tandem for fast and reliable genome-based classification and nomenclature of prokaryotes. Nucleic Acids Research, 50(D1), D801–D807. 10.1093/nar/gkab902

22. Million-Weaver, S., & Camps, M. (2014). Mechanisms of plasmid segregation: Have multicopy plasmids been overlooked? Plasmid, 75, 27–36. 10.1016/j.plasmid.2014.07.002

23. Mistou, M. Y., Sutcliffe, I. C., & Van Sorge, N. M. (2016). Bacterial glycobiology: Rhamnose-containing cell wall polysaccharides in gram-positive bacteria. FEMS Microbiology Reviews, 40(4), 464–479. 10.1093/FEMSRE/FUW006

24. Mohr, K. I. (2018). Diversity of myxobacteria—we only see the tip of the iceberg. In Microorganisms (Vol. 6, Issue 3). MDPI AG. 10.3390/microorganisms6030084

25. Muñoz-Dorado, J., Marcos-Torres, F. J., García-Bravo, E., Moraleda-Muñoz, A., & Pérez, J. (2016). Myxobacteria: Moving, killing, feeding, and surviving together. In Frontiers in Microbiology (Vol. 7, Issue MAY). Frontiers Media S.A. 10.3389/fmicb.2016.00781

26. Nies, D. H. (n.d.). MINI-REVIEW Microbial heavy-metal resistance.

27. Overbeek, R., Olson, R., Pusch, G. D., Olsen, G. J., Davis, J. J., Disz, T., Edwards, R. A., Gerdes, S., Parrello, B., Shukla, M., Vonstein, V., Wattam, A. R., Xia, F., & Stevens, R. (2014). The SEED and the Rapid Annotation of microbial genomes using Subsystems Technology (RAST). Nucleic Acids Research, 42(D1). 10.1093/nar/gkt1226

28. Pal, C., Bengtsson-Palme, J., Kristiansson, E., & Larsson, D. G. J. (2015). Co-occurrence of resistance genes to antibiotics, biocides and metals reveals novel insights into their coselection potential. BMC Genomics, 16(1). 10.1186/s12864-015-2153-5

29. Parker, J. (2001). Bacteria. In Encyclopedia of Genetics (pp. 146–151). Elsevier. 10.1006/rwgn.2001.0102

30. Ramirez, M. S., Traglia, G. M., Lin, D. L., Tran, T., & Tolmasky, M. E. (2014). Plasmid-Mediated Antibiotic Resistance and Virulence in Gram-Negatives: the Klebsiella pneumoniae Paradigm. Microbiology Spectrum, 2(5). 10.1128/microbiolspec.plas-0016-2013

31. Rodríguez-Beltrán, J., DelaFuente, J., León-Sampedro, R., MacLean, R. C., & San Millán, Á. (2021). Beyond horizontal gene transfer: the role of plasmids in bacterial evolution. In Nature Reviews Microbiology (Vol. 19, Issue 6, pp. 347–359). Nature Research. 10.1038/s41579-020-00497-1

32. Schmartz, G. P., Hartung, A., Hirsch, P., Kern, F., Fehlmann, T., Müller, R., & Keller, A. (2022). PLSDB: Advancing a comprehensive database of bacterial plasmids. Nucleic Acids Research, 50(D1), D273–D278. 10.1093/nar/gkab1111

33. Seemann, T. (2014). Prokka: Rapid prokaryotic genome annotation. Bioinformatics, 30(14), 2068–2069. 10.1093/bioinformatics/btu153

34. Sheng, D., Chen, X., Li, Y., Wang, J., Zhuo, L., & Li, Y. (2021). ParC, a New Partitioning Protein, Is Necessary for the Active Form of ParA From Myxococcus pMF1 Plasmid. Frontiers in Microbiology, 11. 10.3389/fmicb.2020.623699

35. Stamatakis, A. (n.d.). RAxML Version 8: A tool for Phylogenetic Analysis and Post-Analysis of Large Phylogenies. http://bioinformatics.oxfordjournals.org/

36. Sun, X., Chen, X. jing, Feng, J., Zhao, J. yi, & Li, Y. zhong. (2011). Characterization of the partitioning system of Myxococcus plasmid pMF1. PLoS ONE, 6(12). 10.1371/journal.pone.0028122

37. Tamura, K., Stecher, G., & Kumar, S. (2021). MEGA11: Molecular Evolutionary Genetics Analysis Version 11. Molecular Biology and Evolution, 38(7), 3022–3027. 10.1093/molbev/msab120

38. Vassallo CN and Wall D. (2019). Self-identity barcodes encoded by six expansive polymorphic toxin families discriminate kin in myxobacteria. PNAS. 116 (49) 24808–24818. 10.1073/pnas.1912556116

39. Wrótniak-Drzewiecka, W., Brzezińska, A. J., Dahm, H., Ingle, A. P., & Rai, M. (2016). Current trends in myxobacteria research. In Annals of Microbiology (Vol. 66, Issue 1, pp. 17–33). Springer Verlag. 10.1007/s13213-015-1104-3

40. Zhao, J. Y., Zhong, L., Shen, M. J., Xia, Z. J., Cheng, Q. X., Sun, X., Zhao, G. P., Li, Y. Z., & Qin, Z. J. (2008). Discovery of the autonomously replicating plasmid pMF1 from Myxococcus fulvus and development of a gene cloning system in Myxococcus xanthus. Applied and Environmental Microbiology, 74(7), 1980–1987. 10.1128/AEM.02143-07

41. Johansen, A. (2021). The functions of microorganisms. Aarhus University. https://envs.au.dk/en/research-areas/microorganisms-in-the-environment/the-functions-of-microorganisms (Johansen,2021)

42. National Center for Biotechnology Information (NCBI)[Internet]. Bethesda (MD): National Library of Medicine (US), National Center for Biotechnology Information; [1988](https://www.ncbi.nlm.nih.gov/)

43. Genome List - Genome - NCBI (nih.gov)

